# Simple in-vitro single stranded linear and circular DNA preparation and validation via SELEX using phosphor-derived modifications

**DOI:** 10.1101/2023.02.11.528153

**Authors:** Seyed Vahid Hamidi, Jonathan Perreault

## Abstract

Interest in preparation of single stranded circular DNA library has been increasing recently, therefore developing a simple and efficient method for circular DNA generation will be very useful for all procedures and techniques that are dependent on single stranded circular DNA preparation. In this study a new simple method for in vitro preparation of circular single stranded DNA is proposed. We hypothesized that using a phosphorylated-phosphorothioated primer would not affect the efficiency of PCR reactions, but, more importantly, would suppress the activity of Lambda Exonuclease enzyme even if it is phosphorylated. The produced phosphorylated single stranded DNA is ready to be circularized via a ligation reaction using a bridging oligonucleotide. Several optimizations and enhancements have been conducted in the ligation reaction, notably by embedding an extra thymine nucleotide at the ligation site to compensate for the additional adenosine nucleotide added by Taq during the PCR reaction. In addition, the performance of the proposed method has been validated by selecting linear and circular aptamers against MERS-CoV spike protein during 15 successive cycles of SELEX. Because this new method is simple and user-friendly, it has a potential to be automated for high-throughput purposes and may further stir growing interests in preparation of single stranded circular DNA and its applications.

## Introduction

Due to the broad applications of in-vitro single stranded DNA (ssDNA) preparation in molecular biology numerous techniques have been developed for this purpose. So far, ssDNA preparation methods have been utilized for systematic evolution of ligands by exponential enrichment (SELEX) (1), oligonucleotide microarray preparation (2), PCR based padlock probe (PLP) generation (3), circular aptamer selection (4–6), single stranded labeled probe production (7), coupled PCR-rolling circle amplification (RCA) for gene production (8). In addition, it has been recently used for linear and circular ssDNA library generation for DNA sequencing (9), production of origami nanostructure (10) as well as long ssDNA donor for genome editing using clustered regularly interspaced short palindromic repeats (CRISPR) (11–14). A good example of the use of ssDNA is aptamers. Aptamers are small ssDNA or RNA molecules that fold into a well-defined three-dimensional structure with a high affinity and specificity for their target molecules. Also, they have better thermal stability, lower cost and are easier to modify compared to antibodies. They are selected by SELEX procedure which starts with a random oligonucleotide library (e.g. 10^15^ sequences) followed by four key steps: (i) specific binding of oligonucleotides (aptamer) with the target, (ii) extracting the bound oligonucleotides, (iii) amplifying of the extracted sequences with PCR (iv) producing an enriched pool of single stranded aptamer sequences that will be used again in step i, and so on. After completion of the SELEX procedure, the candidate aptamers are sequenced for characterization (15). However, developing a simple, fast, efficient and userfriendly technique to produce pure ssDNA is still challenging (16). The majority of ssDNA producing procedures such as denaturing polyacrylamide gel electrophoresis (PAGE) and biotin-streptavidin based techniques are dependent on double strand DNA (dsDNA) denaturation and detachment of the sequence of interest from the complementary strand. Denaturing PAGE has been frequently utilized for pure ssDNA preparation, however it is not convenient for high-throughput approaches because it is complex and time consuming (16–18). In addition, asymmetric PCR and biotin-streptavidin based methods are also criticized because of dsDNA impurity by-products that are generated during the process (7,16,19–21). On the other hand, enzymatic degradation of undesired strand via Lambda exonuclease enzyme is a simple and robust way to produce pure ssDNA. It is reported that this enzyme has 20 folds more activity on 5’-phosphorylated strands of dsDNA than on hydroxylated strands, causing the hydroxylated strands to get single stranded following digestion (19,21,22), but in principle this also precludes its use for any application that require the presence of a 5’-phosphate, such as preparation of PLPs and circular DNA.

The effect of chemically modified nucleotides resistant to nucleases including phosphorothioate bond modifications for ssDNA generation has already been investigated (23,24). In this study, phosphor-derived modifications have been used for in-vitro production of ssDNA, linear or circular depending on the application. For this purpose, PCR was performed using phosphorylated-phosphorothioated forward primer and phosphorylated reverse primer using a 100 bases template (Fig. 1, A and B). In the current work we show that although Lambda exonuclease enzyme has robust digestion activity on phosphorylated strand of dsDNA, its degradation activity is completely suppressed on phosphorothioate modified strand even if it enjoys 5’ phosphorylation modifications (Fig. 1C). In addition, the produced phosphorylated ssDNA can be circularized using complementary strands without prior phosphorylation steps (Fig. 1, D and E). Therefore, the whole procedure of circular ssDNA preparation has become much faster and simpler. In addition, an increased DNA ligation efficiency using T4 DNA ligase has obtained by embedding an additional thymidine nucleotide exactly at the ligation site to account for the adenine added by Taq during PCR. Therefore, this simple, fast and robust method will be useful for specific ssDNA applications and is a good candidate for automation purposes due the broad application of ssDNA preparation in molecular biology. In addition, this technique has been successfully applied on SELEX for linear and circular aptamer selection against MERS-CoV spike protein. This coronavirus is a serious health concern, with close to 30% of affected patients dying from it (25).

**Figure 1.**
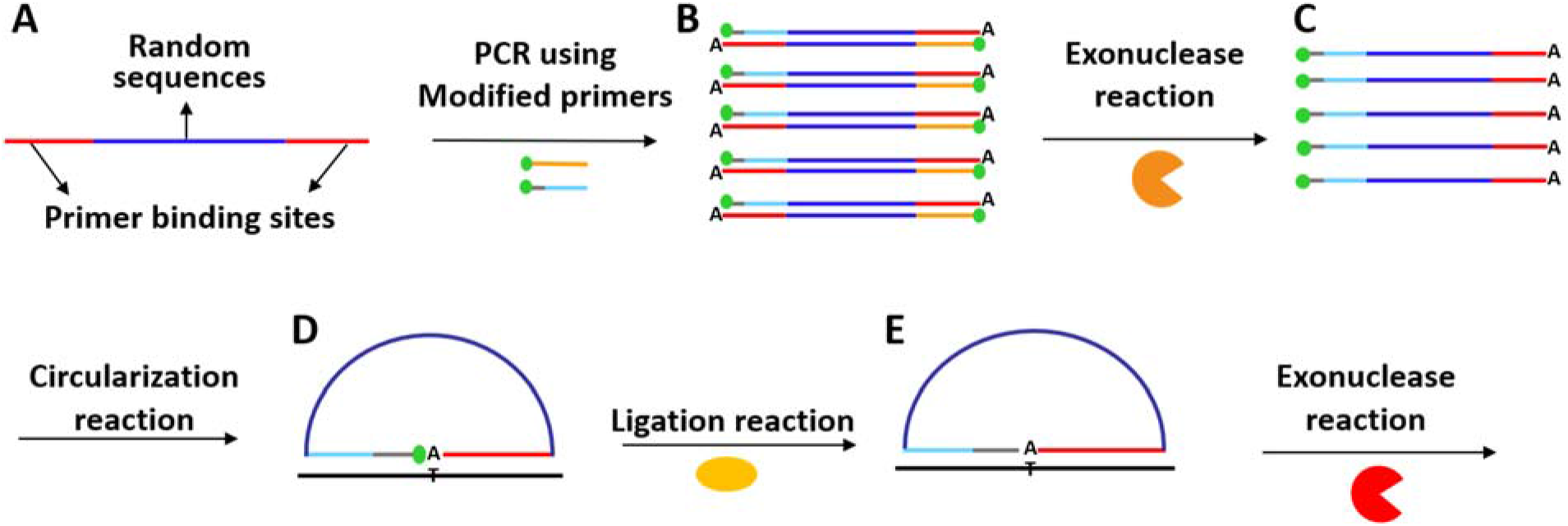
PCR products circularization strategy. (A) Schematic representation of the target used with specific sequences as PCR primers binding sites (red line) at the extremities and random sequences (dark blue line) in the middle. (B) PCR reaction using reverse primer (orange line) with 5’ phosphorylation modification (green dot) and forward primer (light blue line) with 5’ phosphorylation modification (green dot) and five phosphorothioate bonds (gray line) at the far 5’end of the primer. The “A” illustrates the adenine added by the Taq DNA polymerase at the 3’ end of its product. (C) Exonuclease reaction by utilising the Lambda exonuclease enzyme (orange semi-circle) for degradation of non-phosphorothioated reverse strand. (D) ssPCRP circularization via complementary strand (black line). (E) Sealing head to tail of ssPCRP through ligation reaction using a ligase enzyme (yellow ellipse). Finally, non ligated ssPCRP are degraded by the action of Exonuclease I enzyme (red semi-circle)

## Results

### PCR optimization using modified primers

A gradient PCR was performed with normal primers (suggested Tm of 61 °C) using a range of temperatures from 58 to 68 °C to find the best annealing temperature for a sharp PCR band. The sharpness of PCR product bands improved by increasing the annealing temperature and it is at 68 °C that we obtained the best band (data are not shown). Thereafter, PCR using modified primers were performed at 68 °C including with: a phosphorylated reverse primer, as well as phosphorothioated, phosphorylated-phosphorothioated, normal and normal-phosphorylated forward primers (Fig. 2). As it can be seen from Fig. 2, applying phosphorylation and phosphorothioate bond modifications did not have any effects on the quality of PCR bands, including for the PCR reaction using phosphorylated reverse primer and protected-phosphorylated forward primer, the focus of this study. Furthermore, the performance of PCR reactions using modified primers were evaluated by qPCR (Fig. S1). Although signal intensity was decreased by ~25% using modified phosphorylated and phosphorylated/phosphorothioated forward primers, the PCR product quantity is still high enough for all required applications.

**Figure 2.**
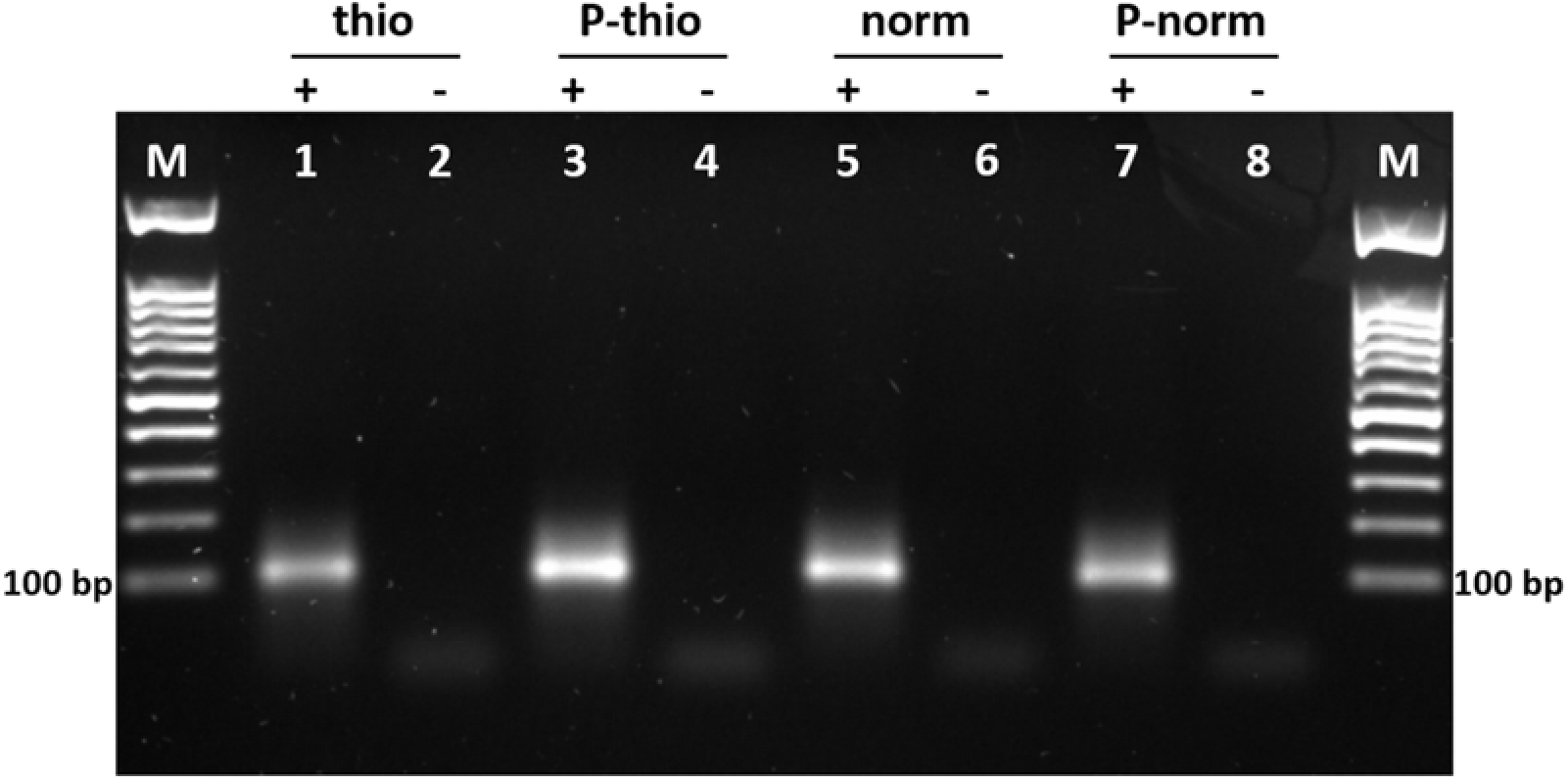
PCR reaction using modified primers. Lanes M: Markers; lanes 1, 3, 5 and 7: positive reactions (with template); lanes 2, 4, 6 and 8: negative reactions (without template). All PCR reactions have been done by utilising a phosphorylated reverse primer (1 μM). Lanes 1 and 2: PCR reaction with phosphorothioated forward primer. Lanes 3 and 4: PCR with phosphorothioated-phosphorylated forward primer. Lanes 5 and 6: PCR products using normal forward primer (no modifications). Lane 7 and 8: PCR with phosphorylated forward primer. All PCRs have been done in the presence of 1 μM of forward primers and 0.02 μM of template.

The quality of PCR products with one distinct length dsDNA is important to get high yield and pure linear ssDNA preparation through exonuclease treatment and subsequently for circular DNA formation (21). Thus, optimization of PCR conditions such as primers and template concentrations, number of cycles and also prudent design of primers to avoid primer dimer formation as well as ensuring specific binding of primers to template is important to avoid pervasive effects on obtaining a defined dsPCR band (21,22).

### Phosphorothioate protects 5’-phosphorylated DNA from degradation

DNA amplification through PCR generates dsDNA products (26). As previously mentioned, the Lambda exonuclease enzyme has an exodeoxyribonuclease activity on dsDNA substrates from 5’ to 3’ and is about 20 times more active on a 5’ phosphorylated terminus compared to a non phosphorylated one. Thus, by phosphorylating one of the primers of the PCR reaction it is possible to use the Lambda exonuclease to easily get a single stranded PCR product (ssPCRP) from a PCR product (22,27,28). Therefore, in this work Lambda exonuclease digestion reactions have been evaluated by employing a phosphorylated reverse primer (to serve as substrate for digestion reactions) and several forward primers with different chemical modifications. All PCR products (size of 100 base pairs) are aligned with the 100 base pair mark of the DNA ladders (Fig. 3A: lanes 2, 5, 8 and 11, as well as lanes M). Whereas ssPCRPs (Fig. 3A: lanes 3, 6 and 9) ran lower in the gel and at the same level as the single stranded 100 bases template (Fig. 3A: lanes 1 and 14) confirming that ssPCRPs became single stranded after Lambda exonuclease treatment. Moreover, we confirmed that the Lambda exonuclease-treated PCR products have been converted to ssDNA, we degraded the DNA with Exonuclease I, a phosphodiesterase enzyme that degrades linear ssDNA in the 3’ to 5’ polarity (29–31). Since all ssPCRPs were digested by Exonuclease I (Fig. 3A: lanes 4, 7 and 10) it confirmed the single stranded nature of ssPCRPs. In addition, because of its 3’ to 5’ exodeoxyribonuclease activity, Exonuclease I it was not blocked by phosphorothioate bonds since these modifications are located at the 5’ extremities of the sense strands (23,24). In principle, using a phosphorylated reverse primer would allow the production of ssPCRP after Lambda exonuclease digestion, however using a phosphorylated forward primer leads to both strand being degraded (Fig. 3A lane 12). To obtain ssPCRP, most laboratories ensure that the forward strand is protected by using a 5’ hydroxyl-forward primer (Fig. 3A: lane 9). Unfortunately, this is not compatible with applications that require a phosphorylated 5’end. Although PCR sense strands without any modifications and with phosphorothioate bonds modification were protected from degradation (Fig. 3A: lanes 6 and 9), it is noticeable that the phosphorylated sense strand with five phosphorothioate-modified bonds was protected from digestion by Lambda enzyme (Fig. 3A: lane 6 and Fig. 1C). Lambda exonuclease action inhibition by phosphorylated-phosphorothioated modifications enables us to circularize the ssPCRP using a complementary strand without requiring to phosphorylate ssPCRP after exonuclease reaction. Thus, in addition to excluding the phosphorylation step after exonuclease action, with this approach the protected phosphorous group can be used for ssPCRP circularization by ligase enzymes (32,33). Therefore, the whole circularization procedure has become much simpler and more efficient.

**Figure 3.**
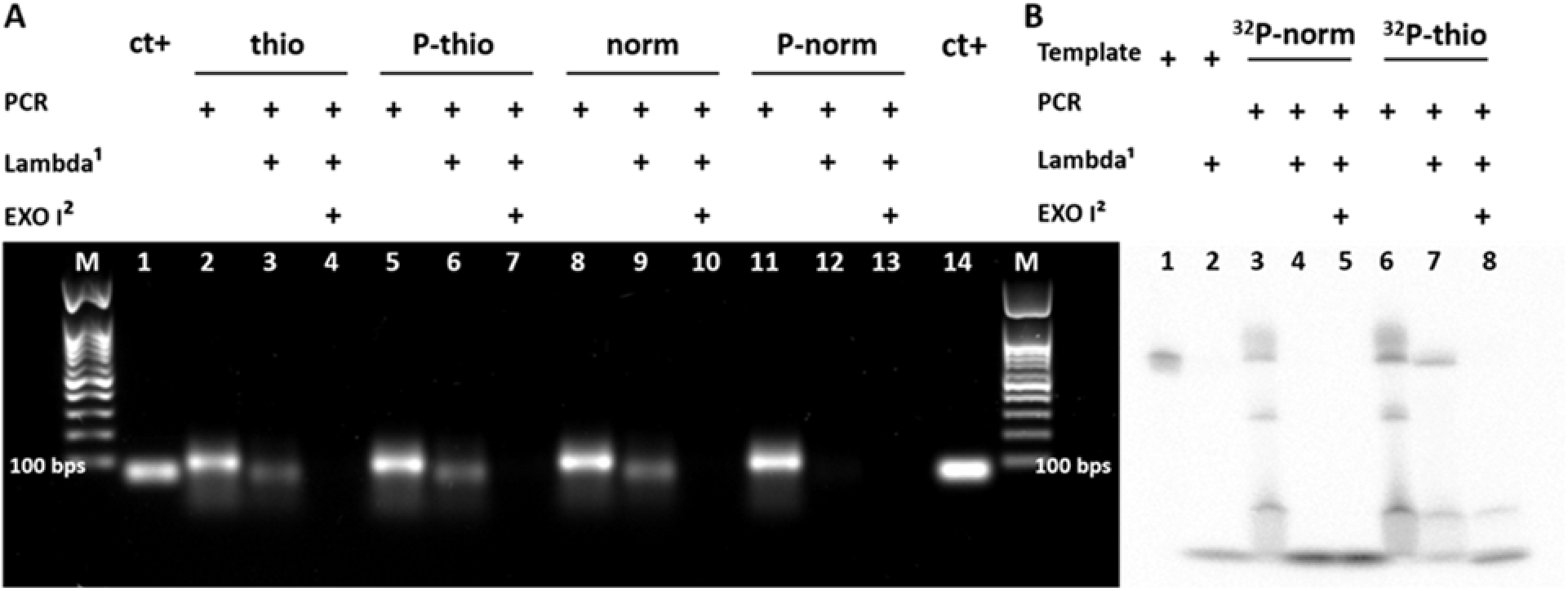
Preparation of ssPCRP. (A) Visualization via a 2% agarose gel. Lanes M: Markers. Lanes 1 and 14: single stranded target with final length of 100 bp (80 μM). Lanes 2, 5, 8 and 11: PCR products. Lanes 3, 6, 9 and 12: ssPCRP preparation with Lambda Exonuclease enzyme from PCR products. Lanes 4, 7, 10 and 13: ssPCRP treatment using Exonuclease I enzyme. All PCR reactions were performed using phosphorylated reverse primers (1 μM). Lanes 2, 3 and 4; 5, 6 and 7; 8, 9 and 10; and 11, 12 and 13 have been done using phosphorothioated, phosphorothioated-phosphorylated, normal and phosphorylated forward primers, as indicated in the figure, in final concentration of 1 μM, respectively. (B) ssPCRP analysis using 6% denaturing PAGE. Lanes 1 and 2: ^32^P-labeled 100 bases oligonucleotide (0.1 pmol), intact or treated with Lambda exonuclease, respectively. Lanes 3 and 6: PCR reaction with phosphorylated reverse primers (1 μM). Lanes 4 and 7: ssPCRP preparation by Lambda Exonuclease. Lanes 5 and 8: ssPCRP incubation with Exonuclease I enzyme. Lanes 3, 4 and 5; and lanes 6, 7 and 8: PCR reactions were done by employing normal-^32^P-labeled and phosphorothioated-^32^P-labeled forward primers, respectively, in final amount of 4 pmoles.

In order to verify the inhibition of Lambda exonuclease action at the very 5’phosphorylated end of PCR strands modified by phosphorothioated bonds, highly sensitive [γ-32P] radio-labeling was used. Phosphor imaging of Lambda exonuclease degradation pattern of [γ-32P] radio-labeled PCR products was evaluated using a phosphorylated reverse primer and [γ-32P] radio-labeled forward primers without and with phosphorothioate modifications (Fig. 3B: lanes 3 and 6, respectively) and employing denaturing 6% PAGE (Fig. 3B). The two different undigested PCR products showed similar band patterns, with the main PCR bands migrating to the same level as the 100 bases template (Fig. 3B: lanes 3, 6 and 1). Importantly, the sense normal PCR strand was completely digested by Lambda Exonuclease (Fig. 3B: lane 4), whereas the sense phosphorothioated PCR strand remained intact during degradation reaction (Fig. 3B: lane 7). As opposed to agarose gels (Fig. 3A), double stranded PCR products and ssPCRP run similarly on denaturing PAGE because hydrogen bonds between bases are weakened in the latter (34,35). Afterward, radio-labeled-phosphorothioated ssPCRP was again digested by Exonuclease I enzyme to confirm the single stranded property of phosphorothioated ssPCRP (Fig. 3B: lane 8). Finally, the [γ-32P] nucleotide residues resulting from digestion accumulated at the bottom of the gel, further verifying the digestion activity of exonuclease enzymes.

### ssPCRP circularization

Circular DNA has some advantages over linear DNA, including higher stability against misfolding and exonucleases as well as compatibility to be utilized as template for exponential signal generation (5,36,37). In order to convert linear ssPCRP to circular ssPCRP, six different complementary strands in different sizes including 50, 42 and 35 bases were employed as bridging oligonucleotides for ligation (Fig. 4A). Taq DNA polymerase enzymes add an additional adenosine nucleotide at the 3’ ends of PCR strands during amplification reaction (38–40). Thus, one additional thymine (T) nucleotide was embedded in the complementary strands exactly at the ligation spot to compare the ligation efficacy with thymine-less counterparts. Furthermore, in an attempt to improve the efficiency of ligation, two different DNA ligase enzymes were used: the T4 DNA ligase and the thermostable Taq DNA ligase enzymes (Fig. 4, B and C). The ligation efficiency was affected by the presence of an additional T, the length of complementary strand and the type of ligase enzyme. Since covalently closed circular ssPCRPs have lower migration rates as compared with linear ssPCRPs in agarose or PAGE gels (3,41), we used gel electrophoresis to monitor ligation efficacy. As previously mentioned the overall size of ssPCRP is 100 bases thus if a bridging oligonucleotide of 50 complementary bases is utilized for ssPCRP circularization, it hybridizes with half the length of ssPCRP which likely hinders the full hybridization due to the lack of flexibility of the allowed by a ssDNA region the same length of the dsDNA region (Fig. 4A, 1). However, by decreasing the complementary strand sizes, less bases in ssPCRP are occupied in base-pairing which favors more efficient hybridization reactions (Fig. 4A, 1 and 2). When T4 DNA ligase and complementary strands with an additional T are used for ligation reactions, elevated ligation efficiency was seen for 42 and 35 bases complementary strands compared to 50 bases for ssPCRP circularization (Fig. 4B lanes 1, 2, 4 and 6). However, when complementary strands without additional T were used, we witnessed lower ligation efficacy even in the presence of 42 and 35 bases strands (Fig. 4B lanes 3 and 5), reducing by ~30% the amount of circularized DNA; and, perhaps more importantly, increasing by as much as ~7 fold the amount of linear DNA left over (from 5% to 37% for 35 bp bridging oligonucleotide). This can be explained by the adenosine nucleotide added by Taq DNA polymerase enzyme which leads to a one base mismatch pairing at the ligation site. Therefore, in absence of the added T to the bridging oligonucleotide, T4 DNA ligase is not able to properly seal head to tail of the ssPCRP efficiently because of the base mismatch at this strategic spot (33,42,43). On the other hand, when ligation is done via the Taq DNA ligase enzyme, satisfying ligation bands were detected just by employing 35 bases complementary strands even without an additional T (Fig. 4C lanes 5 and 6). Better ligation efficiency was expected via Taq DNA ligase because of its thermostability and ability to apply several denaturing-annealing cycles for ligation reaction (44). Although we observed lower selectivity for Taq ligase than T4 ligase enzyme at the ligation spot, for the purpose of circularization of a library this does not represent a significant problem. Indeed, by using four T nucleotides at the ligation junction, we likely increased the ability of the DNA strand to be ligated even when using only four instead of the ideal five T nucleotides complementary to the extra A nucleotide added during PCR at the ligation junction (Table S1). Therefore, we tried ligation efficiency in presence of different bases at the ligation site (Fig. 4, E and F). Better ligation efficiencies were obtained for both T4 and Taq DNA ligase enzymes when bridging strands with an additional T are used. In addition, better signal intensity was observed by hyperbranched RCA (HRCA) reaction on circularized ssPCRP using + T bridging strand which shows higher ssPCRP circularization rates compared to the strand without the additional T (Fig. S2). All in all, T4 DNA ligase would be a better candidate for ssPCRPs circularization in this study due to its efficiency (90%-95% of DNA ligated with bridging oligonucleotide of 42 and 35 bp), due to the fact that it works at moderate temperature and also due to its lower price that can eventually improve the throughput of approaches that use the proposed method; overall providing a more user-friendly technique (45).

**Figure 4.**
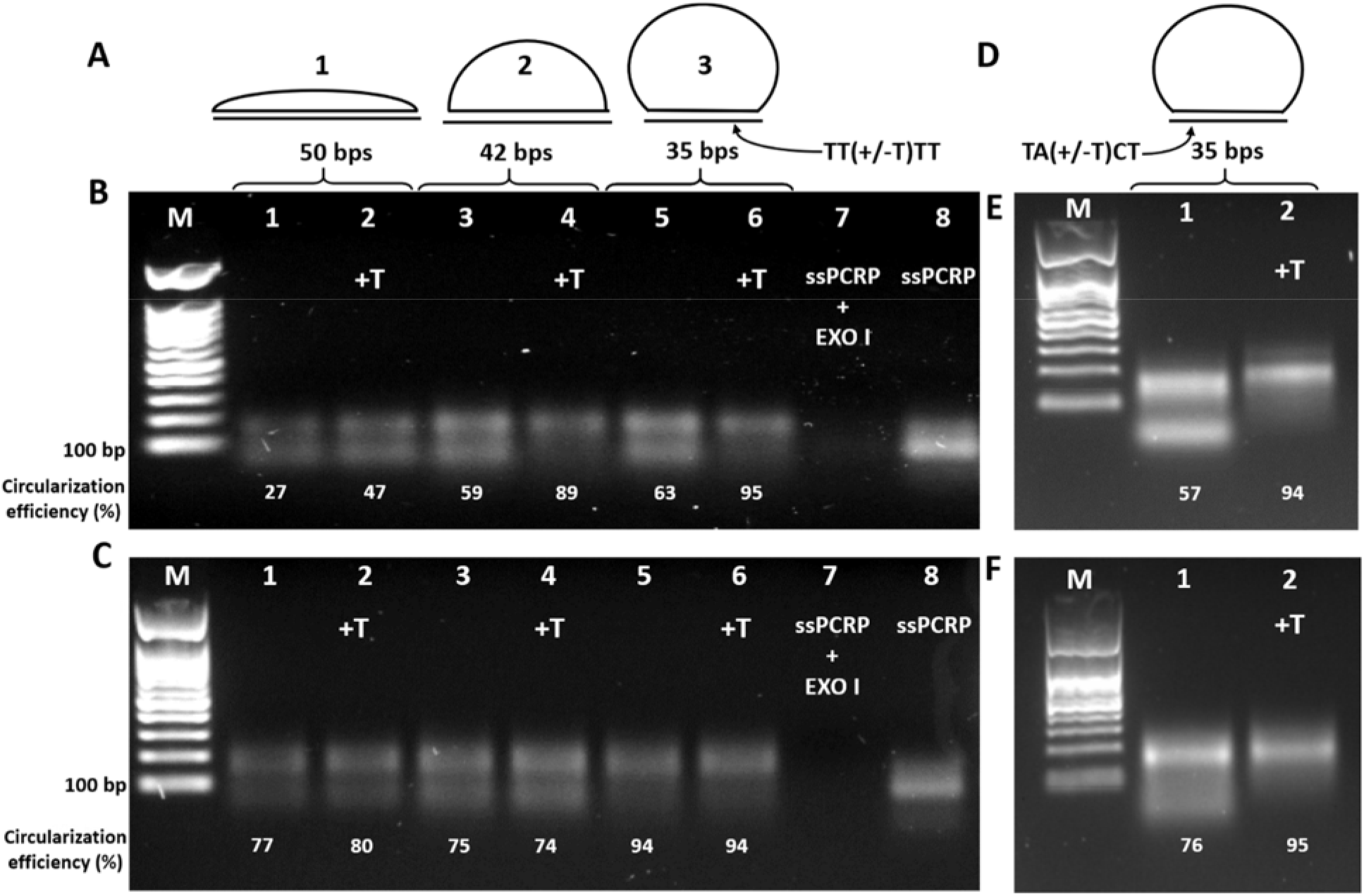
ssPCRP circularization using complementary strands. (A) Schematic representation of ssPCRP circularization with 50, 42 and 35 complementary bases bridging oligonucleotide containing TT(+/-T)TT at ligation site. (B) ssPCRP circularization with T4 DNA ligase. Lane M: Marker. Lanes 1, 3 and 5: circularization reaction in presence of 50, 42 and 35 bp complementary strands (TTTT sequences at ligation sites) and lanes 2, 4 and 6 in combination with 50, 42 and 35 bp complementary strands with an additional T nucleotide at the ligation site (TTTTT) at a final concentration of 1 μM, respectively. Lane 7: ssPCRP treatment with Exonuclease I. Lane 8: ssPCRP. (C) ssPCRP head to tail sealing using Taq DNA ligase enzyme. Order of the lanes are the same as part A. (D) Schematic illustration of ssPCRP circularization using 35 bases complementary strand containing specific sequences at ligation site. (E and F) ssPCRP circularization via T4 and Taq DNA ligase enzymes respectively. Lane M: Marker; Lanes 1: Circularization reaction using bridging oligonucleotides containing TACT sequence at ligation position; Lanes 2: Circularization reaction via additional T in ligation position.

### HRCA reaction using circularized ssPCRPs

To confirm that ssPCRP is covalently closed though a ligation reaction, circular ssPCRP has been used as a circular DNA template in presence of PCR primers (forward and reverse) and highly processive Phi29 DNA polymerase for exponential HRCA amplification. Signal amplification caused by HRCA reaction leads to production of a large amount of dsDNA which in turn can be monitored in real-time by an intercalating fluorescent molecule (SYBR green) (46). Fluorescence intensity increases overtime during exponential HRCA and it almost reaches a plateau after 120 minutes where the amplification reaction gets saturated (Fig. 5A). Moreover, the intensity of fluorescence signal is proportional to the initial concentration of circular ssPCRP because lower amplification signal slopes are observed by two folds diluting of circular ssPCRP (Fig. 5A, from top to bottom). The product of real-time HRCA reaction was migrated on agarose gel to confirm production of high molecular weight and hyperbranched nature of the generated DNA, which leads to long smeared DNA bands (Fig. 5B: lanes 1 to 8: from high to low concentrations).

**Figure 5.**
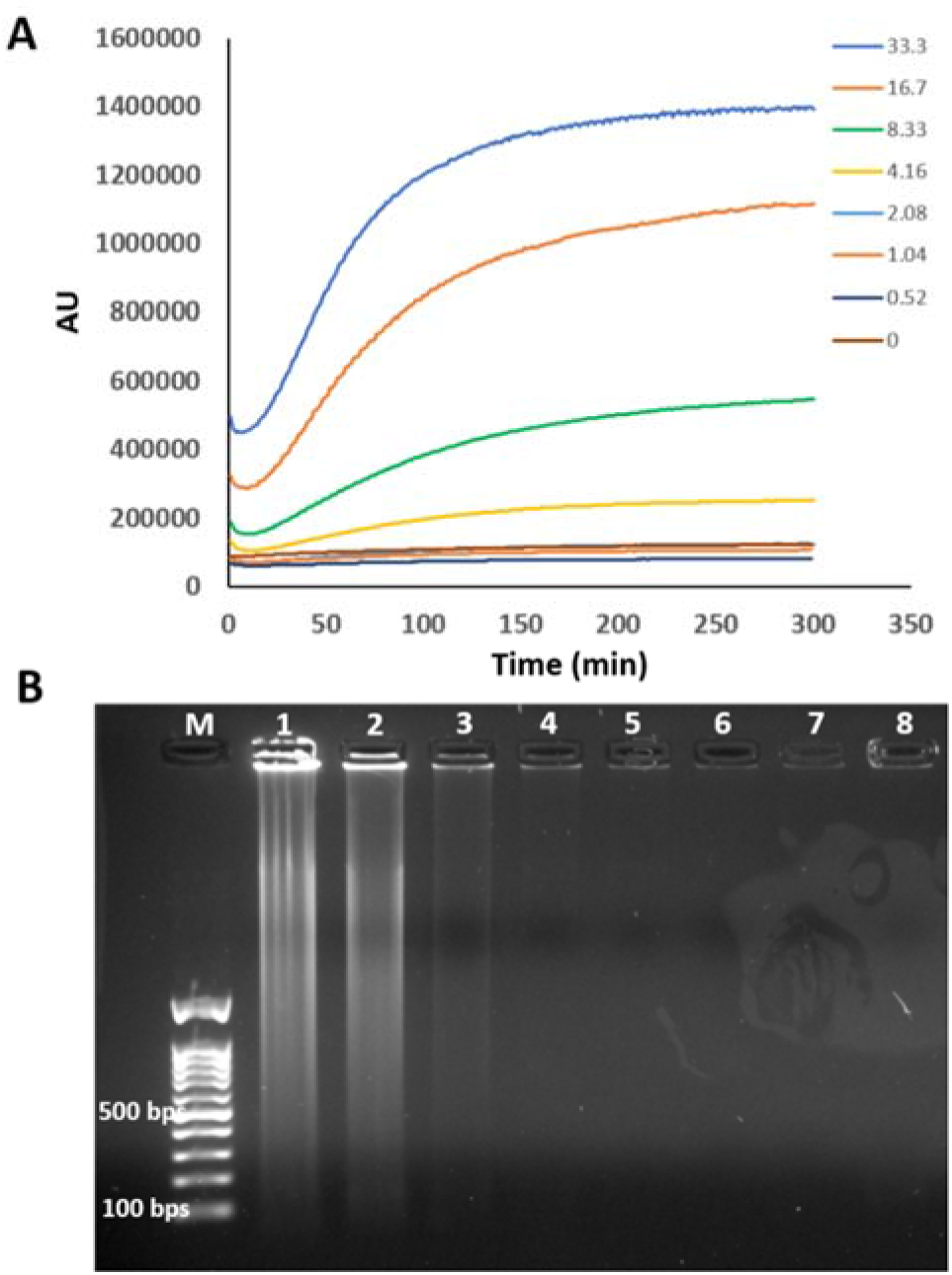
HRCA reaction using circular ssPCRP. (A) Real-time monitoring of HRCA reaction via real-time PCR machine in different concentration of circular ssPCRP (from top to bottom: 33.3, 16.7, 8.33, 4.16, 2.08, 1.04, 0.52 and 0 nM). (B) Visualization of HRCA product by agarose gel electrophoresis. Lane M: Marker. Lanes 1 to 7: HRCA product from the highest concentration of circular ssPCRP (33.3 nM) to the lowest one (0.52 nM). Lane 8: Negative control (no ssPCRP).

### SELEX based on phosphorylated-phosphorothioate primers

In order to verify the performance of the proposed method in a relevant application, this technique has been used to select circular and linear aptamers through 15 successive rounds of SELEX against the whole MERS-CoV spike protein as well as a fragment from spike protein. In addition, four negative selections were performed using agarose magnetic beads and lysozyme protein immobilized on agarose magnetic beads to exclude nonspecific aptamers from the library (Fig. S3). By increasing SELEX cycles, the amount of ligand and ligand/library incubation time were gradually decreased and washing times and number of washing were gradually increased. Therefore, based on the utilized conditions, selected aptamers through generation 15 are supposed to have lowest *K*_D_ compared to other cycles. In Table 1, selected aptamers were categorized based on their cluster sizes obtained from generation 15 and frequency of mutations that appeared during SELEX cycles. These results confirm the applicability of the proposed technique to generate diverse libraries for the SELEX procedure. After 15 successive cycles of SELEX, the first rank selected aptamers from all four libraries including circular and linear aptamers against full size spike protein and RBD fragment were characterized for *K*_D_ determination. To do this, aptamers labeled with Cy5 were used in order to be able to check *K*_D_ using powerful NanoTemper Monolith machine. In addition, since the circular aptamers are covalently closed circular DNA and thus have no ends for 5’ Cy5 modifications, a short labeled complementary strand which is specific to the 5’ end region of aptamers were used (Fig. 6A-upper). The same approach was employed for linear aptamers *K*_D_ determination to be able to compare affinities with circular aptamers (Fig. 6A-below). As it can be seen from Fig. 6B, an average *K*_D_ of ~ 50 μM was obtained using Labeled complementary strand and Monolith strategy. In addition, the selectivity of selected aptamers was also determined in the presence of bovine serum albumin (BSA) as a non-specific ligand and no specific bindings were detected (data are not shown). In addition, since labeled short oligo strategy was used in this experiment, the selectivity of procedure was determined in the presence of full spike and RBD and labeled complementary stands (without adding aptamers) and no affinity was observed (data are not shown). This shows that the complementary strand has not affinity to the specific targets.

**Table 1.**
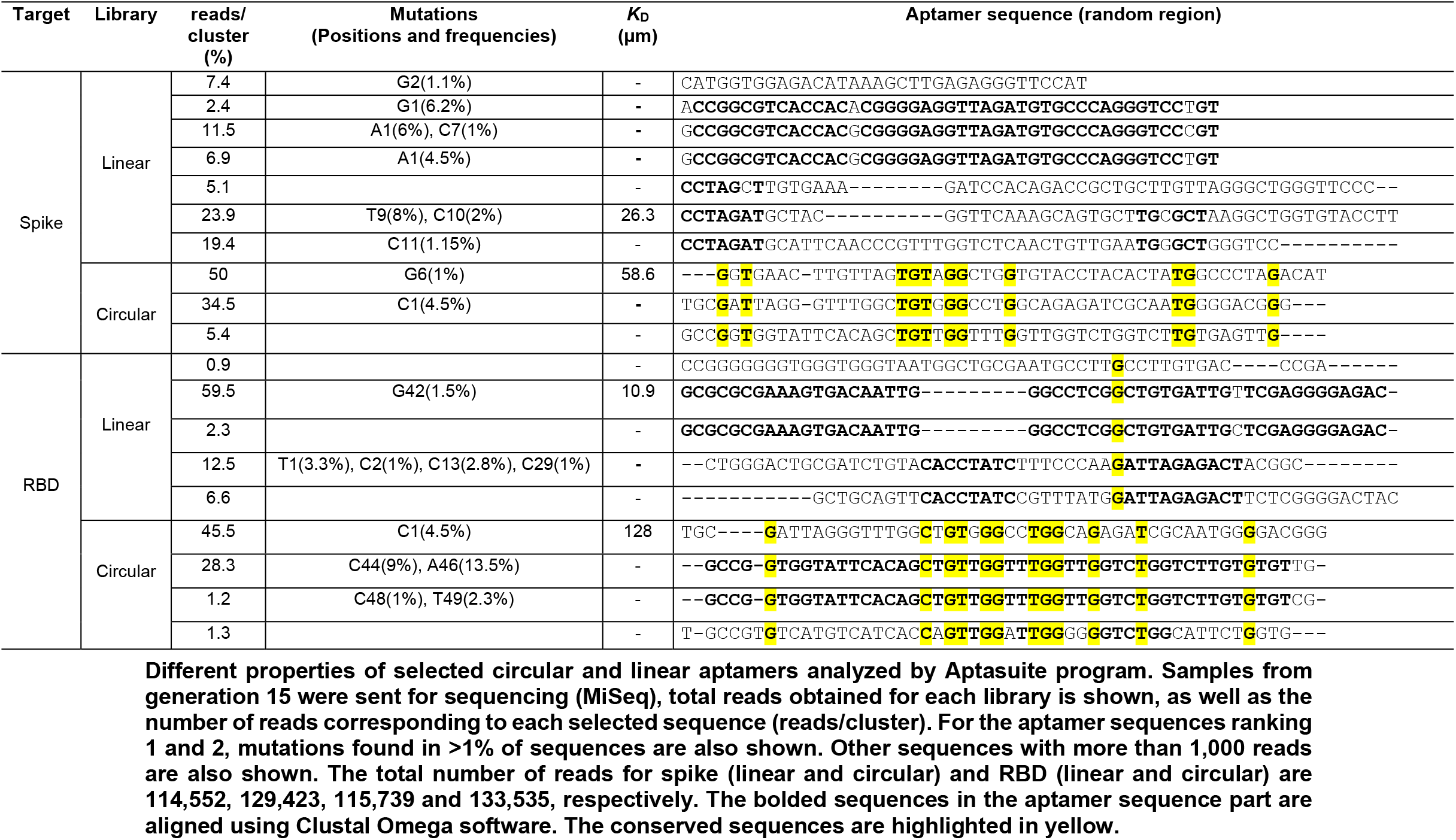
Selected circular and linear aptamers.

**Figure 6.**
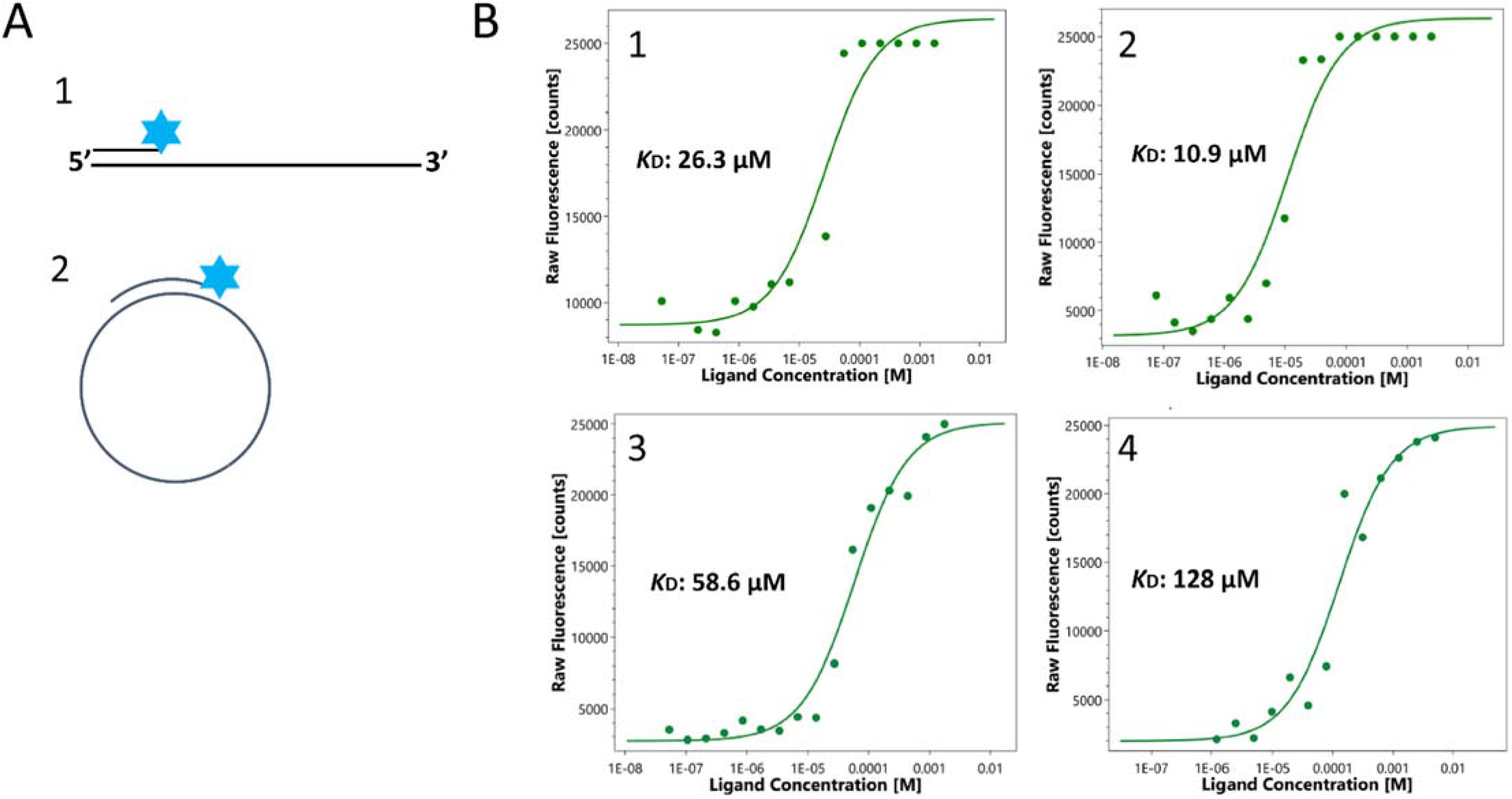
*K*_D_ determination. (A) Schematic illustration of selected linear and circular hybridization using labeled Cy5 complementary strand. (B) Binding affinity of selected aptamers (Linear/full spike (1), Linear/RBD (2), circular/full spike (3) and circular/RBD (4)) in the presence of 5 nM of labeled aptamers and 1mM to ~ 10^-8^ M of ligands (full spike and RBD).

## Discussion

As compared with other reported in vitro ssDNA generating techniques in the literature, the proposed method is more user-friendly and has improvements as well as other advantages (Table 2) (3,8,16,22,47–49). This method is much simpler and faster than conventional gel-based ssDNA preparation because the whole reaction happens in just a tube which bypasses the time consuming steps of gel preparation and running, as well as DNA elution and precipitation that are followed in gel-based procedures (18). In addition, ssDNA generation through biotin-streptavidin and NaOH treatment causes possible disruption of hydrogen bonds and hydrophobic forces between streptavidin and biotin which leads to detachment of biotinylated strand and rehybridization with complementary strands in the solution, and thus presence of unwanted dsDNA in the ssDNA preparation (19,20). The same dsDNA impurity also happens in asymmetric PCR reaction where some level of DNA complementary to the strand of interest remains (7,16). Lambda exonuclease digestion of a phosphorylated reverse strand bypasses the problems mentioned above (22), but is not compatible with applications that require a phosphate group at the 5’end of the DNA strand. However, embedding chemical phosphorothioate modifications at the 5’end inhibits the Lambda exonuclease enzyme activity leaving the strand intact after the degradation of the reverse strand (19,27). Therefore, in this study we devised a 5’ phosphorylated strand that is not affected by the digestion reaction by also including phosphorothioate modifications. Our approach allows the omission of the normally required phosphorylation step before ssPCRP circularization, thus making the whole procedure simpler (4). Furthermore, in comparison with circle-to-circle amplification (C2CA) which is another enzymatical way for single stranded linear or circular DNA production (50–52), the proposed method is faster and less complicated and no dsDNA impurity is produced during the process. Indeed, in C2CA in order to reach the sequence with desire polarity it is necessary to do at least two rounds of C2CA reaction and repeat even more rounds to get more products (51). C2CA also as more dsDNA impurities because it includes steps of polymerisation and endonuclease reactions where the produced ssDNA is partially double stranded (50).

**Table 2.** Comparison of different aspects of reported methods in the literature with the proposed technique.

| Method | Application | Type of modifications | Amplification Method(s) | Gel based technique | dsDNA impurity | circDNA<sup>1</sup> preparation | Reference |
| --- | --- | --- | --- | --- | --- | --- | --- |
| Lambda Exo<sup>2</sup> | ssDNA preparation | Phosphorothioate & phosphorylation | PCR | - | - | + | Present work |
|  | ssDNA preparation | Phosphorylation | PCR | - | - | - | (22) |
| Biotin-streptavidin | SNP<sup>3</sup> detection | Biotinylation | PCR | - | + | + | (3) |
|  | Genomic DNA production | Biotinylation & phosphorylation | PCR & RCA | - | + | + | (8) |
|  | ssDNA preparation | Biotinylation | PCR | + | + | - | (47) |
| C2CA<sup>4</sup> | Gene detection | - | RCA | - | + | + | (51) |
| Denaturing PAGE | Aptamer selection | HEGL<sup>5</sup> & fluorescein | PCR | + | - | - | (49) |
| Asymmetric PCR | ssDNA preparation | Phosphorylation | PCR | - | + | -<sup>6</sup> | (16) |

<sup>1</sup>Circular DNA
<sup>2</sup>Lambda exonuclease enzyme
<sup>3</sup>Single nucleotide polymorphism
<sup>4</sup>Circle to circle amplification
<sup>5</sup>Hexaethyleneglycol
<sup>6</sup>This study has not performed circularization with the ssDNA, but it would theoretically be possible

By employing a template with random sequences and non-proof-reading Taq DNA polymerase enzyme as well as DNA circularization optimization using a complementary strands and DNA ligase enzymes, we confirmed the applicability of this method for either linear or circular aptamer selection through the SELEX procedure. Furthermore, because the method does not rely on gels and the whole reactions are done in a tube, it is possible to automate ssDNA preparation using this procedure. This technique is a good alternative for coupling of PCR with HRCA for exponential gene generation and for PCR-based PadLock Probes, as well as single stranded library production in general (2,3,7–9). Sequencing technologies such as Nanopore and PacBio use circularized DNA templates to increase quality of sequencing (53–55), the method explained herein could provide an alternative to the current protocols which could, in addition, also allow ligation of a few DNA fragments before circularization, thus providing a longer fragment for sequencing of even longer reads compared to the current methodology. Moreover, it was recently found that long ssDNA provides a very efficient way of making CRISPR-mediated knock-ins (11), in this case also, the inclusion of a phosphate at the 5’ end of PCR products allows ligation of different ssPCRP to make ssDNA beyond the typical range of PCR products, allowing insertions of larger DNA fragments in the genome, in addition to simplifying the procedure for ssDNA preparation (or reducing the cost if Megamers are used). Overall, the proposed method is likely to help many technical advancements in a number of areas in molecular biology and genetics.

## Experimental procedures

### Targets, primers and aptamers

The 100 bases template with two primer binding sites (25 bases for each) and 50 random bases in the middle is provided form Integrated DNA Technologies (IDT). All primers (forward and reverse) either with modifications (phosphorothioate bonds and phosphorylation) or without modifications and aptemrs were purchased from Sigma Aldrich Company. Also, complementary strand targets were obtained from IDT and Alpha ADN company (Quebec, Canada) (Table S1).

### PCR and ssPCRP ligation reactions

PCR amplification was performed in PCR reaction mixture including target (0.02 μM), phosphorylated reverse primer (1 μM), phosphorothioated-phosphorylated forward primer (1 μM), 0.20 μM deoxynucleoside triphosphate (dNTP) (DGel Electrosystem, Canada), milli-Q water, 10X HotStarTaq buffer (QIAGEN) and 0.5 units HotStarTaq DNA Polymerase (QIAGEN) in final volume of 50 μL. PCR was achieved in 12 cycles and at optimized melting temperature (tm) of 68.2 °C thorough C1000 touch thermal cycler (Bio-rad). For single stranded PCR product (ssPCRP) preparation, each 10 μL of double stranded PCR products were treated with 5 units Lambda exonuclease enzyme (New England Biolabs, NEB) at 37 °C for 45 min which is followed by 10 min at 75 °C for enzyme inactivation.

To circularize ssPCRP, ligation reactions with T4 DNA ligase were prepared: 1 μM of ssPCRP was added to 1 μM complementary strands in 1X T4 ligation buffer (NEB) in a final volume of 10 μL. For the circularization reaction, DNA was denatured by increasing temperature to 95 °C for 5 min and then gradually cooled down to room temperature for DNA hybridization and circular ssPCRP formation. Afterward, the ligation reaction was performed by introducing 10 units T4 DNA ligase enzyme (NEB, USA) and incubating at 20 °C for 1 hour to seal the 5’ and 3’ ends of the ssPCRP. Alternatively, it is possible to use the Taq DNA ligase (NEB) with ~20 cycles of denaturation at 95 °C for 1 min followed by annealing and ligation steps using proper Tm for 1 min (Tm of 68, 64 and 60 °C for 50, 42 or 35 complementary bases between strands, respectively). After ligation, either by using T4 or Taq DNA ligase enzymes, 10 units Exonuclease I enzyme (NEB) is added to ligation mixture for non-circular ssPCRP degradation.

### HRCA reaction

HRCA reaction was performed using different dilutions of circular ssPCRP (33.33, 16.66, 8.33, 4.16, 2.08, 1.04, 0.52 and 0 nM) in a final volume of 30 μL of amplification mixture including: 1X Phi29 DNA polymerase buffer (NEB), 0.3 μM non-modified PCR forward and reverse primers, 0.40 μM dNTP, milli-Q water, SYBR green 1X (Invitrogen), 10 μg bovine serum albumin (BSA) and 8 units Phi29 DNA polymerase enzyme (NEB). Amplification reactions were monitored for 5 hours at 30 °C in MicroAmp fast reaction tubes with cap (Applied Biosystems) and using real-time PCR machine (Applied Biosystems).

### Agarose gel electrophoresis

DNA illustration by agarose gel electrophoresis has been conducted by exploiting 2% agarose gel and 1X TAE buffer (1 mM EDTA and 40 mM Tris-acetate, Fisher Scientific) and by applying constant voltage of 120 V for 40 to 60 min. Thereafter, gels were illuminated under UV radiation via gel doc (Bio-Rad) and 1X pre-added Gel Stain (TransGen Biotech). Power supply and DNA ladder were purchased from Bio-Rad and Bio-Helix, respectively.

### [γ-32P] radio-labeling

[γ-32P] labelling of 100 bases target and forward primers (phosphorothioated and non-phosphorothioated) were manipulated in a final volume of 20 μL labeling mixture containing 1X T4 polynucleotide kinase (PNK) enzyme buffer (NEB), milli-Q water, 20 pmoles of DNA substrate, 5 μCi [γ-32P] ATP (PerkinElmer) and 10 units T4 PNK enzyme (NEB) at 37 °C for 30 min.

### Visualization via polyacrylamide gel electrophoresis (PAGE)

Phosphor imaging of [γ-32P] radio-labelled DNA sequences was accomplished through 6% denaturing PAGE constituted of 1X TBE (89 mM Tris base, 2 mM EDTA and 89 mM boric acid, Fisher Scientific) (57), 8 M urea (BioShop, Canada), 6% Acrylamide/Bis-acrylamide (BioShop, Canada), 16 μL N,N,N’,N’-tetramethylethane-1,2-diamine (TEMED) (Fisher Scientific), 300 μL ammonium persulfate (APS) 10% (BioShop, Canada) in a total volume of 40 mL. Denaturing PAGE ran for 1 hour under constant power of 15 watt (W), subsequently the gel was exposed to a phosphorimager cassette (GE Healthcare) for 10 min and finally the cassette scanned by a Typhoon FLA 9500 biomolecular imager (GE Healthcare).

### Recombinant MERS-CoV spike protein

Recombinant MERS-CoV spike protein and MERS-CoV spike protein fragment were produced from the MERS-CoV genome sequences (human betacoronavirus 2c EMC/2012) which encodes spike protein and receptor binding domain (RBD), respectively. For construction of MERS-CoV Spike/RBD Protein fragment (RBD, aa 367-606), the DNA sequence that encodes MERS-CoV RBD protein (#AFS88936.1, Glu367-Tyr606) was fused to the C-terminus of the Fc region of rabbit IgG. RBD obtains 464 aa and has molecular mass of 51.5 kDa with purity of > 80 % that is determined by SDS-PAGE. For construction of MERS-CoV Spike Protein (S1+S2 ECD, aa 1-1297, His Tag), the DNA sequence that is encoding the MERS-CoV spike protein extracellular domain (AFS88936.1, Met1-Trp1297) was fused with a polyhistidine tag at the C-terminus. The recombinant spike protein MERS-CoV extracellular domain obtains 1291 aa and has molecular mass of 142.52 kDa with purity of > 85 % as determined by SDS-PAGE. For expression of recombinant MERS-CoV spike proteins Baculovirus insect cells were used. The recombinant RBD and full spike proteins were directly purchased from Sino Biological, China.

### Spike proteins immobilization

Recombinant spike proteins immobilization on agarose magnetic beads (CNBr-activated SepFastTM MAG 4HF, BioToolomics, UK) was performed based on the protocol provided by BioToolomics. In this work, 10 mg of each recombinant protein was immobilized on 300 mg/ml agarose magnetic beads. After doing several steps for spike protein functionalization on agarose magnetic beads, the immobilized beads washed several times using SELEX buffer and magnet to remove un-bound spike proteins from stock solution. Thereafter immobilized spike protein was diluted in 500 μl of SELEX buffer which was used as stock spike protein solution to perform 15 cycles of SELEX. This solution should be vortexed well before addition to each cycle to make sure that homogeneous concentration of immobilized protein is added to SELEX cycles.

### SELEX procedure

Circular library preparation was performed in final T4 ligation mixture of 100 μl including 1X T4 DNA ligase buffer (NEB, USA), 1 μM phosphorylated library, 1 μM 35 bp complementary strand, 20 U T4 DNA ligase enzyme (NEB, USA) and milliQ water for 1 hour at 16 °C which was followed by enzyme inactivation for 20 mins at 80 °C. Thereafter, the ligation reaction was treated by 10 U of Exonuclease I enzymes (NEB, USA) for 1 hour at 37 °C for degradation of non-circularized library. Then, the exonuclease was inactivated at 80 °C for 20 mins.

The first round of SELEX was started separately in the presence of 100 pmol of linear and circular libraries using immobilized spike proteins and spike proteins fragments. During the first round the linear and circular libraries were incubated for 2 hours with immobilized spike proteins on agarose magnetic beads at room temperature in 1 ml of 1× SELEX buffer (20 mM Tris-HCl (pH:7.4), 2 mM CaCl2, 5 mM KCl, 100 mM NaCl, 1 mM MgCl2). Thereafter, the bound libraries were collected by applying a magnet (Invitrogen, USA) which is followed by supernatant removal and washing agarose beads with 500 μL 1× SELEX buffer. Afterward the beads were diluted in 20 μL of SELEX buffer and 5 μL of the diluted library was directly used for enrichment step using PCR protocol described in the main text of paper and in the final volume of 50 μL. In this step a phosphorothioated forward primer and phosphorylated reverse primer were used for the linear library and a phosphorothioated/phosphorylated forward primer and phosphorylated reverse primer utilized for the circular library. After verification of PCR bands using agarose gel, ssPCRPs were produced from PCR products using the previously described procedure and at this point the linear library has been regenerated for another round of SELEX. For the circular library the produced phosphorylated libraries were circulated using the process explained earlier with a T4 DNA ligase enzyme for sealing the library head to tail. The positive selection process has been done for 15 rounds of SELEX using circular and linear libraries and using the spike protein and the spike protein fragment as targets. Moreover, four SELEX rounds were performed as negative selection cycles using non-specific targets including agarose magnetic beads and lysozyme protein immobilized on agarose magnetic beads (Table S2). From SELEX rounds 1 to 15 concentration of ligands and incubation time have gradually decreased, while number of washes as well as washing time have gradually increased. The detailed information about SELEX conditions used for each SELEX round is shown in Table S2.

### KD determination using Monolith device

5’ phosphorylated circular aptamers against full spike/RBD were circularized using procedure mentioned in section 4.5.2, however in this ligation reaction 35 bp complementary strand without was used since the aptamers were directly purchased and were not the product of PCR (ssPCRP). Thereafter, selected linear/circular aptamers were equally diluted with 5’ Cy5 labeled complementary strand to reach the optimum concentrations of 5 nM where satisfying fluorescence intensity was obtained for *K*_D_ determination using Monolith device (NanoTemper, Germany). Affinity of the selected aptamers was evaluated in 16 test tubes where spike/RBD proteins were diluted by double from the initial concentration 1 mM and then 5 nM of aptamer/Cy5 labeled complementary strand complex were added to each protein dilution in final volume of 20 μl. The labeled aptamer/protein mixture is incubated for 30 min at room temperature before loading them in Monolith NT.115 Capillary (NanoTemper, Germany) and determining binding affinity using Monolith machine.

### qPCR reaction

Real-time monitoring of qPCR reaction using modified primers including normal, phosphorylated, phosphorothioated and phosphorylated/phosphorothioated forward primers as well as a phosphorylated reverse primer were done in qPCR reaction mixtures including 1X reaction mixture of GoTaq 2-Step RT-qPCR kit (Promega, USA), 400 nM forward and reverse primers and in final volume of 20 μL. PCR was performed a total of 20 cycles and at an annealing temperature of 68 °C and via an Applied Biosystems qPCR machine.

## Funding and additional information

SV Hamidi received fellowships from Armand-Frappier foundation and Fonds québécois de la recherche sur la nature et les technologies (FRQNT). JP is a junior 2 FRQS research scholar. Other funding, including for open access charge: NSERC [RGPIN-2019-06403].

## Author contributions

Conceptualization, S.V.H.; Resources, J.P.; Methodology, S.V.H.; Validation, S.V.H. and J.P.; Data analysis, S.V.H. and J.P.; Experimental work: S.V.H.; Writing—Original Draft Preparation, S.V.H.; Writing—Review and Editing, S.V.H. and J.P.; Supervision: J.P.; Funding Acquisition, J.P.

