## Supplementary materials for "Simple in-vitro single stranded linear and circular DNA preparation and validation via SELEX using phosphor-derived modifications"

Table S1	Oligonucleotide sequences used in this study.

| PCR and RCA forward primers | 5′-AACGCCTATGTCTCGTTGAAGCAGC3′ |
| --- | --- |
|  | 5′-P-AACGCCTATGTCTCGTTGAAGCAGC-3′ |
|  | 5′-A*A*C*G*C*CTATGTCTCGTTGAAGCAGC-3′ |
|  | 5′-P-A*A*C*G*C*CTATGTCTCGTTGAAGCAGC-3′ |
|  | 5′-P-T*A*C*G*C*CTATGTCTCGTTGAAGCAGC-3′ |
| PCR and RCA reverse primer | 5′-P-TTGCCGGTGACAGACTGCTTGCATA-3′ |
|  | 5′-P-CTGCCGGTGACAGACTGCTTGCATA-3′ |
| Template/library | 5′-AACGCCTATGTCTCGTTGAAGCAGCNNNNNNNNNNNNNNNNNNNNNNNNNNNNNNNNNNNNNNNNNNNNNNNNNNTATGCAAGCAGTCTGTCACCGGCAA-3′ |
|  | 5′-P-AACGCCTATGTCTCGTTGAAGCAGCNNNNNNNNNNNNNNNNNNNNNNNNNNNNNNNNNNNNNNNNNNNNNNNNNNTATGCAAGCAGTCTGTCACCGGCAA-3′ |
| Complementary strands | 5′-GCTGCTTCAACGAGACATAGGCGTTTTGCCGGTGACAGACTGCTTGCATA-3′ |
|  | 5′-GCTTCAACGAGACATAGGCGTTTTGCCGGTGACAGACTGCTT-3′ |
|  | 5′-CAACGAGACATAGGCGTTTTGCCGGTGACAGACTG-3′ |
|  | 5′-CAACGAGACATAGGCGTACTGCCGGTGACAGACTG-3′ |
| Complementary strands with additional T | 5′-GCTGCTTCAACGAGACATAGGCGTTTTTGCCGGTGACAGACTGCTTGCATA-3′ |
|  | 5′-GCTTCAACGAGACATAGGCGTTTTTGCCGGTGACAGACTGCTT-3′ |
|  | 5′-CAACGAGACATAGGCGTTTTTGCCGGTGACAGACTG-3′ |
|  | 5′-CAACGAGACATAGGCGTATCTGCCGGTGACAGACTG-3′ |
| Labeled complementary strand | Cy5-ACATAGGCGTT |

**P indicates phosphorylation modifications**

***shows phosphorothioate bond modifications**

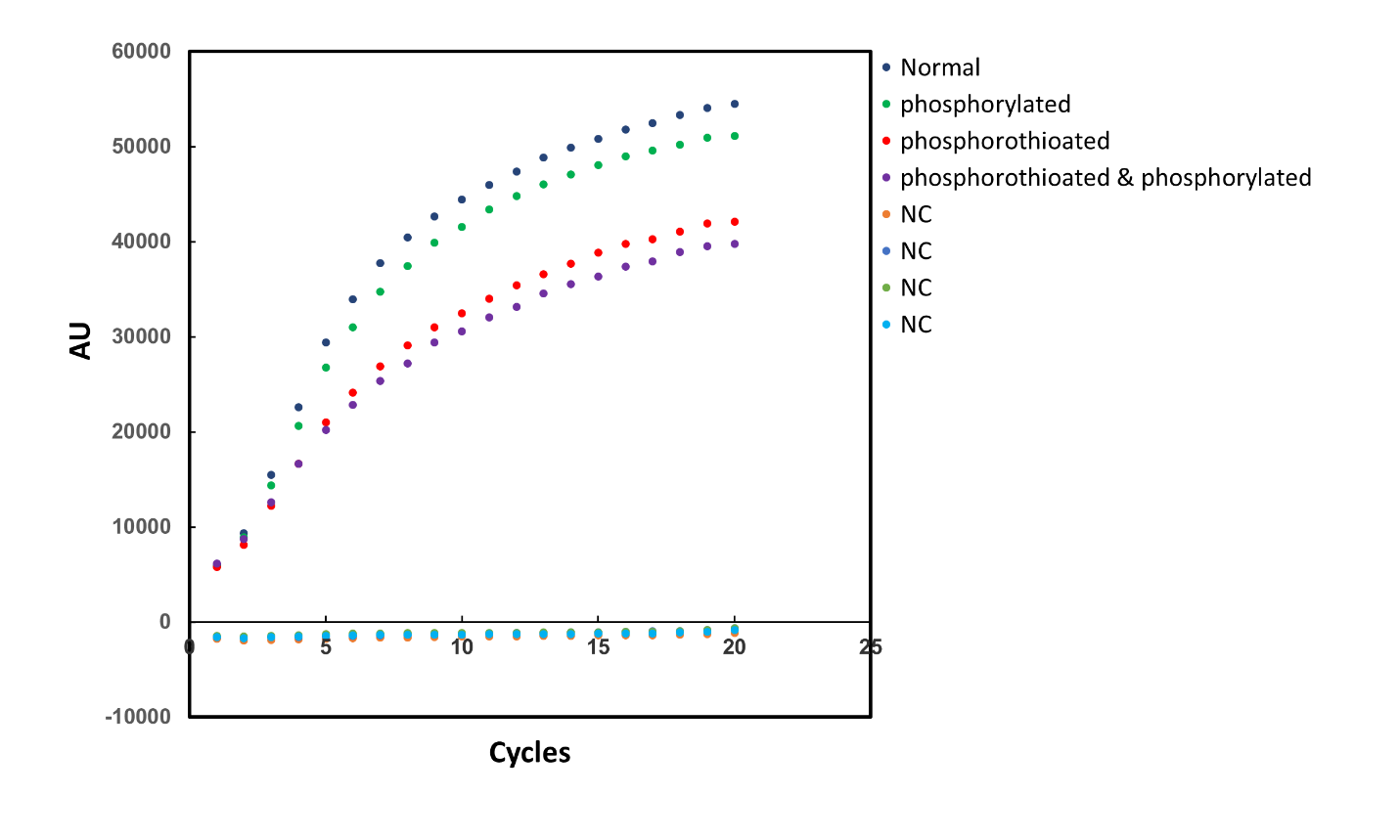

Figure S1	Real-time monitoring of modified PCR primers using qPCR technique. Primers used are as indicated in the legend within the figure, NC are the corresponding negative controls.

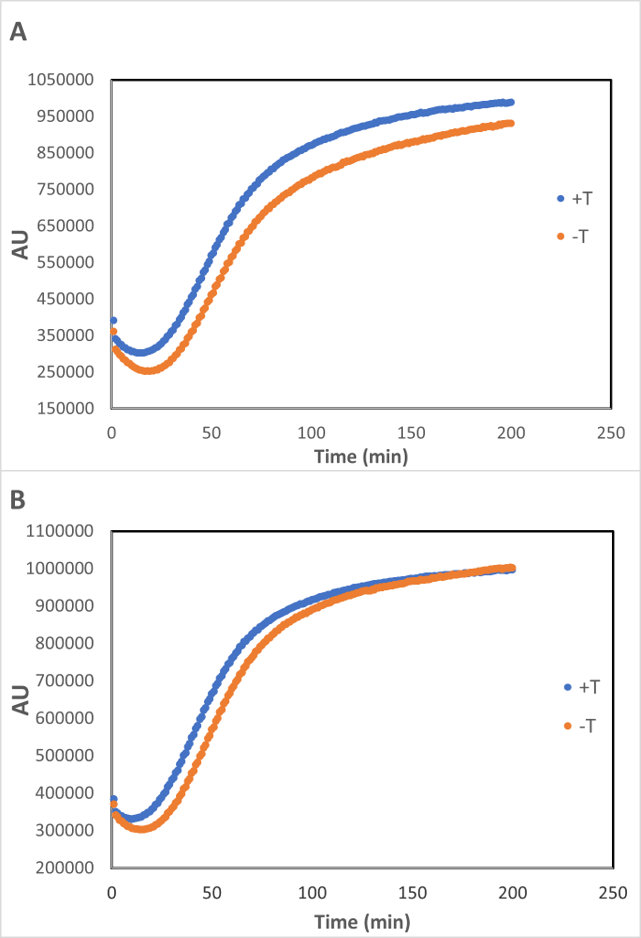

Figure S2	HRCA reaction using circularized ssPCRP with and without additional T. (A) Monitoring of HRCA reaction via circularized ssPCRP sealed by T4 DNA ligase using specific complementary strand. (B) HRCA reaction through circular ssPCRP sealed by Taq DNA ligase and specific complementary strand.

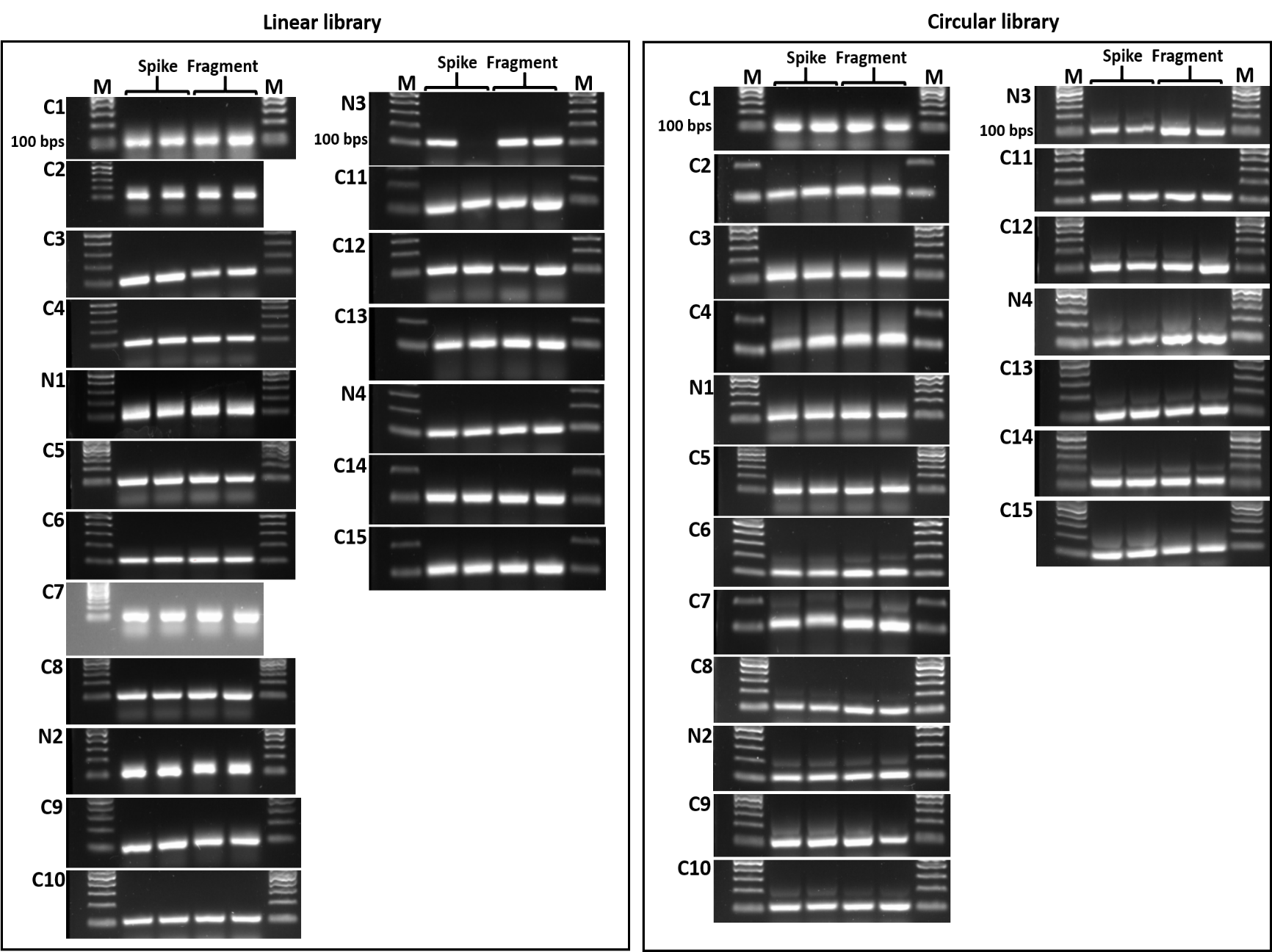

Figure S3	Amplification of selected libraries after each cycle of SELEX for linear and circular aptamer selection targeting MERS-CoV spike protein. M, C and N refer to ladder, SELEX cycle, and negative selection respectively. The reason why additional bands are seen in PCR amplification products of circular libraries, especially generations 6, 7, N2 is that it is likely two ssPCRs were hybridized with two complementary strands during ligation process and thus dimers were produced during ssPCRs circularization reaction. PCR products are shown in duplicate for each full spike or RBD because during each SELEX round one PCR product was used for the next SELEX round and the other one was kept as a backup in case something went wrong during the SELEX process, both are shown.

Table S2	Detailed conditions used for circular and linear aptamer selection against MERS CoV spike protein.

| Target | Library | Rounds | Incubation time (min) | No of washes | Added target  (µl) | Washing time  (min) |
| --- | --- | --- | --- | --- | --- | --- |
| Full spike  & fragment (RBD) | Linear & circular | C1 | 120 | 1 | 200 | 30 |
|  |  | C2 | 120 | 1 | 20 | 30 |
|  |  | C3 | 120 | 1 | 20 | 30 |
|  |  | C4 | 120 | 1 | 20 | 30 |
|  |  | N1 | 30 | - | 10 (beads) | - |
|  |  | C5 | 105 | 1 | 20 | 30 |
|  |  | C6 | 90 | 1 | 20 | 30 |
|  |  | C7 | 75 | 1 | 20 | 45 |
|  |  | C8 | 60 | 1 | 20 | 45 |
|  |  | N2 | 60 | - | 15 (beads) | - |
|  |  | C9 | 45 | 2 | 20 | 45 |
|  |  | C10 | 30 | 2 | 15 | 45 |
|  |  | N3 | 30 | - | 10 (lysozyme) | - |
|  |  | C11 | 30 | 3 | 15 | 60 |
|  |  | C12 | 30 | 2 | 10 | 60 |
|  |  | C13 | 30 | 3 | 10 | 60 |
|  |  | N4 | 60 | - | 15 (lysozyme) | - |
|  |  | C14 | 10 | 3 | 10 | 60 |
|  |  | C15 | 5 | 2 | 5 | 60 |
